# Extracellular vesicles from retinal pigment epithelial cells expressing R345W-Fibulin-3 induce epithelial-mesenchymal transition in recipient cells

**DOI:** 10.1101/2020.10.05.327221

**Authors:** Mi Zhou, Yuanjun Zhao, Sarah R. Weber, Han Chen, Michael Ford, Matthew T. Swulius, Alistair J. Barber, Stephanie L. Grillo, Jeffrey M. Sundstrom

**Affiliations:** Department of Ophthalmology, Penn State Hershey College of Medicine, Hershey, PA, USA; TEM, Microscopy Imaging Facility, Penn State Hershey Medical Center, Hershey, PA, USA; MS Bioworks LLC, Ann Arbor, MI, USA; Department of Biochemistry and Molecular Biology, Penn State Hershey College of Medicine, Hershey, PA, USA

## Abstract

**Purpose:** Previous studies in our lab found that expression of R345W-Fibulin-3 induces retinal pigment epithelial (RPE) cells to undergo epithelial-mesenchymal transition (EMT). The purpose of the current study was to investigate the size, cargo and function of extracellular vesicles (EVs) derived from RPE cells expressing wild-type (WT)-Fibulin-3 compared to RPE cells expressing the R345W-Fibulin-3 mutation, and to determine the role of these EVs in regulating RPE cell dysfunction.

**Methods:** ARPE-19 cells were infected with luciferase-tagged wild-type Fibulin-3 (WT)- or luciferase-tagged R345W-Fibulin-3 (mutant) using lentivirus. EVs were isolated from the media of ARPE-19 cells by conventional ultracentrifugation or density gradient ultracentrifugation. Transmission electron microscopy (TEM) and cryogenic electron microscopy (cryo-EM) were performed to study the morphology of the EVs. The amount and size distribution of EVs were determined by Nanoparticle Tracking Analysis (NTA). EV protein concentrations were quantified using the DCTM Protein Assay (Bio-Rad). EV cargo were analyzed by unbiased proteomics using LC-MS/MS with subsequent pathway analysis (Advaita). The EV-associated transforming growth factor beta 1 (TGF-β1) protein was measured by Enzyme-linked immunosorbent assay (ELISA). The EV transplant study was conducted and migration ability was evaluated in ARPE-19 cells with or without exposure to EVs by conducting scratch assays.

**Results:** TEM imaging revealed concave-appearing vesicles, and cryo-EM imaging showed spherical vesicles with two subpopulations of EVs: a small group with diameters around 30nm and a large group with diameters around 100nm. Imaging also indicated a greater number of small EVs (~30 nm) in the mutant group compared to the WT group. This result was further confirmed by NTA showing that, in the mutant group, the particle size distributions were smaller than those of the WT EVs. There were no significant differences in EV protein concentrations per EV between WT and mutant groups. Proteomic studies showed that EVs derived from ARPE-19 cells expressing WT-Fibulin-3 contain critical members of sonic hedgehog signaling (SHH) signaling and ciliary tip components, whereas EVs derived from RPE cells expressing R345W-Fibulin-3 contain EMT mediators, including TGF-β-induced protein (TGFBI), vimentin, and mothers against decapentaplegic homolog 4 (SMAD4), indicating that the EV cargo reflects the phenotypic status of their parental cells. Subsequent studies revealed enhanced activity of TGF-β1 associated with mutant EVs compared to WT EVs. Critically, EV transplant studies showed that treatment of recipient RPE cells with mutant RPE cell-derived EVs was sufficient to induce an enhanced migration ability and elevated EMT marker expression of RPE cells.

**Conclusions:** The expression of R345W-Fibulin-3 alters the size, cargo and autocrine function of EVs. Notably, EVs derived from RPE cells expressing R345W-Fibulin-3 are sufficient to induce EMT in uninfected RPE cells.

## INTRODUCTION

Extracellular vesicles (EVs) play a critical role in cell-cell communication and modulate cellular differentiation (Yuyama et al., 2008;Alvarez-Erviti et al., 2011;Weber et al., 2020). In numerous tissues, EVs contribute to the regulation of epithelial-mesenchymal transition (EMT), including in the lungs, breasts, liver, and brain (Vella, 2014;Kim et al., 2016;Chen et al., 2017;van de Vlekkert et al., 2019). For example, myofibroblast-derived EVs are sufficient to induce normal fibroblasts to become myofibroblasts that possess mesenchymal features by upregulating transforming growth factor beta (TGF-β) pathways and EMT drivers (van de Vlekkert et al., 2019). Recent studies have shown that alterations in EV size and cargo are dependent upon the secretion mechanisms and phenotypic status of their parental cells (Zhang et al., 2018). This study examined the role of EV’s secreted from retinal pigment epithelial (RPE) cells in the induction of EMT.

RPE form a monolayer of highly polarized cells that lies posterior to the neuroretina, and are essential for maintaining the health and function of the adjacent photoreceptors. Loss of terminal differentiation and acquisition of a mesenchymal cell phenotype in RPE cells has been shown to occur in degenerative retinal diseases (Ghosh et al., 2018;Goldberg et al., 2018;Wu et al., 2019;Zhou et al., 2020a;Zhou et al., 2020b). As RPE cells become atrophic, the hallmark clinical lesions of sub-RPE lipoprotein deposits and pigmentary changes appear within the macula (Weber et al., 1994;Stone et al., 1999;Vincent et al., 2012).While RPE cells have been shown to secrete EVs (Biasutto et al., 2013;Klingeborn et al., 2017;Atienzar-Aroca et al., 2018;Shah et al., 2018), the role of EVs in RPE function and dysfunction remains to be determined.

An inherited macular degeneration in which RPE dysfunction plays a significant role is Doyne honeycomb macular dystrophy, which results from a single arginine-to-tryptophan point mutation, R345W, in Fibulin-3 (Marmorstein, 2004;Narendran et al., 2005). Fibulin-3 is an extracellular matrix protein that contains six epidermal growth factor (EGF)-like domains followed by a fibulin domain (Zhang and Marmorstein, 2010). The R345W mutation causes Fibulin-3 misfolding, poor Fibulin-3 secretion, and activation of the unfolded protein response (UPR) (Marmorstein et al., 2002;Hulleman and Kelly, 2015). Previously we found that overexpression of R345W-Fibulin-3 in RPE cells activates the UPR, attenuates RPE differentiation and induces EMT in RPE cells (Zhou et al., 2020b).

In this study, we investigated the mechanism by which EVs regulate EMT in RPE cells. EVs were isolated from media of cultured ARPE-19 cells expressing either WT or mutant Fibulin-3. Expression of the mutant protein altered the size of EV’s and upregulated EMT markers including expression of transforming growth factor beta (TGF-β). They also enhanced the migration ability of RPE cells, indicating that EVs can act in a, autocrine manner to influence adjacent RPE cell differentiation. The data show that RPE-derived EV’s strongly influence the phenotype of neighboring cells and likely play an important role in diseases of the posterior eye.

## METHODS

### ARPE-19 Cell Culture

ARPE-19 Tet-On cell with Lentiviral GLuc-tagged WT- or R345W-Fibulin-3 were described previously (Hulleman et al., 2013;Zhou et al., 2020b). Inserted genes have inducible expression in the presence of doxycycline (1ug/ml, Dox, #D9891, MilliporeSigma, Burlington, MA, USA).

### Sample Preparation and Proteomic Analysis

A total of 20 μg of protein was processed using sodium dodecyl sulfate-polyacrylamide gel electrophoresis (SDS-PAGE) with a 4-12% Bis-Tris Mini-gel (Invitrogen) and the 3-(N-morpholino) propanesulfonic acid (MOPS) buffer system. Following electrophoresis, the gel lanes were excised into 40 equal-sized segments and processed by in-gel digestion with trypsin (DigiLab). Briefly, fragments were washed with 25 mM ammonium bicarbonate followed by acetonitrile. Samples were reduced with 10 mM dithiothreitol at 60°C followed by alkylation with 50mM iodoacetamide at room temperature. Digestion was performed with Sequencing Grade Trypsin (Promega, Madison, WI, USA) at 37°C for 4 h. Samples were quenched with formic acid, and the supernatant was analyzed directly without further processing.

Half of each gel digest was analyzed by nano LC-MS/MS with a Waters NanoAcquity high performance liquid chromatography system interfaced to a ThermoFisher Q Exactive mass spectrometer. Peptides were loaded on a trapping column and eluted over a 75 μm analytical column at 350 nL/min. Both columns were packed with Luna C18 resin (Phenomenex). The mass spectrometer was operated in data-dependent mode, with the Orbitrap operating at 60,000 and 17,500 full widths at half maximum (FWHM) for MS and MS/MS, respectively. The 15 most abundant ions were selected for MS/MS. The mass spectra were processed using Mascot set to search the SwissProt Human database (forward and reverse appended with common contaminant proteins) set with Trypsin/P as the enzyme. The resultant Mascot DAT files were parsed into Scaffold (Proteome software) for validation, filtering, and creation of non-redundant protein lists for samples. Data were filtered using a 1% protein and peptide false discovery rate (FDR) and requiring at least two unique peptides per protein.

### Bioinformatics

Proteomic analyses were performed using Advaita iPathway Guide. The log differential expression (Log2DE) was calculated by determining the differential expression of R345W and WT proteins, then taking the log base 2 of that value. Bos Taurus proteins (from bovine serum) and uncharacterized proteins were removed from the analysis.

### Extracellular Vesicle Isolation

Cell culture media (serum-free) from ARPE-19 was filtered through 0.22-um filter and concentrated with Centricon Plus 70 (100K NMWL, MilliporeSigma). EVs for proteomic study were isolated using a sucrose cushion and iodixanol buoyant density gradient ultracentrifugation (Dong-Sic Choi, 2016;Zhou et al., 2020c). Briefly, the concentrated cell culture media was added to the 0.8 M sucrose and 2 M sucrose cushion. After ultracentrifugation, EVs were located at the interface between two sucrose cushion buffer layers. The interface solution was collected and placed at the bottom of an iodixanol buffer. After ultracentrifugation, 10 fractions remained in the tube, with EVs in the third fraction. The 10 fractions were collected and ultracentrifugation was conducted on each. The 10 pellets were re-suspended in 4-(2-hydroxyethyl)-1-piperazineethanesulfonic acid (HEPES)-Buffered Saline (HBS).

EVs for characterization and western blot studies were isolated using differential centrifugation. Briefly, cell culture media was filtered using a 0.22-μm filter and then was concentrated to 1/10th of its original volume using Amicon Centrifugal Filter Ultra-15mL 3K. Concentrated cell culture media was ultracentrifuged at 100,000 x g for 17 h and was washed using HBS at 100,000 x g for 5 h. EV pellets were re-suspended in HBS buffer.

### Nanoparticle Tracking Analysis

EV suspensions were diluted to 1 mL (1:50 to 1:1000) with particle-free water. Each sample was loaded by syringe pump into the NanoSight NS300 (Malvern Instruments Ltd, Malvern, Worcestershire, UK) set in scatter mode, and five 60-second videos were generated. The size distribution and concentration of particles were analyzed and images were acquired using NanoSight software version 3.2 (Malvern Instruments Ltd).

### EV Transplant and Cell Migration Assay

Cell migration assays were performed as described previously (Zhou et al., 2020b). Briefly, ARPE-19 cells were cultured in 96-well ImageLock Microplates (Essen Bioscience Inc., Ann Arbor, MI, USA) to confluence (n=8). A 96-pin WoundMaker™ (Essen Bioscience Inc., Ann Arbor, MI, USA) were used to make scratches. The wells were then washed with PBS to remove cell debris. EVs were added to the cell culture media. EV concentrations (1 x10^10^ vesicles/mL) were chosen according to a previously published study (van de Vlekkert et al., 2019). Wound images were acquired automatically and data were analyzed by the IncuCyte™ software system (Essen Bioscience Inc., Ann Arbor, MI, USA).

### Real-Time PCR

Real-time (RT)-PCR was performed as described previously (Zhou et al., 2020b). In brief, total RNA was extracted from samples using the AllPrep DNA/RNA/Protein Mini Kit (80004, Qiagen Sciences, Inc.) and 0.25 μg of total RNA was reverse transcribed by SuperScript™ IV First-Strand cDNA Synthesis System (18091050, Thermo Fisher Scientific) with a mixture of oligo dT and random hexmer. Amplification of cDNA was performed with optimized PCR primers designed by Primer3 software (https://bioinfo.ut.ee/primer3/) and synthesized by Integrated DNA Technologies (Coralville, Iowa) in SYBR Green master (Rox) system (04913850001, Roche, Basel, Switzerland) on QuantStudio™ 3 Real-Time PCR Systems (ThermoFisher). Detailed information on PCR primers used in this study are provided in Supplemental Table 1. The relative quantity of mRNA was normalized to glyceraldehyde 3-phosphate dehydrogenase (GAPDH) using the comparative 2^-ΔΔCt^ method.

### TGF-β1 Enzyme-Linked Immunosorbent Assays (ELISAs)

Cell culture media was collected from ARPE-19 cells expressing either WT-Fibulin-3 or R345W-Fibulin-3. EVs from media were isolated by using conventional ultracentrifugation and supernatant was also collected. TGF-β1 ELISAs (Quantikine; R&D Systems, Minneapolis, MN, USA) were performed on media, supernatant and EVs from both groups according to kit instructions. Optical densities were determined within 30 minutes with a SpectraMax 190 microplate reader (Molecular Devices, San Jose, CA, USA) at 450 nm with wavelength correction at 570 nm. All TGF-β1 ELISA experiments were run in technical duplicates and biological triplicates.

### Western Blot Analysis

Protein lysates were quantified using the DC™ Protein Assay (Bio-Rad). Total protein (10 μg) was loaded for each sample. Western blots were performed as described previously (Zhao et al., 2018). The primary antibodies used in this study were rabbit anti-GLuc 1:2,000 (E8023S, New England Biolabs), rabbit anti-epidermal growth factor receptor (EGFR), 1:1,000 (sc-03, Santa Cruz Biotechnology), rabbit anti-ALG-2-interacting protein X (Alix) 1:1,000 (ABC40, MilliporeSigma), and mouse anti-Flotilin-1 1:1,000 (610821, BD Biosciences).

### Transmission Electron Microscopy (TEM)

Negative staining was conducted as following. Briefly, the resuspended pellet were fixed in 2% paraformaldehyde. The fixed vesicles (10 ul) were deposited on Formvar carboncoated TEM grids and incubated for 20 minutes. The grids were then washed with PBS, incubated with glutaraldehyde, and washed with water. The vesicles were then stained with uranyl acetate solution and air dried. Vesicles were observed using the JEM1400 TEM (JEOL Ltd., Tokyo, Japan).

### Cryogenic Electron Microscopy (Cryo-EM)

4 μl of EV extract was pipetted onto a freshly glow-discharged Quantifoil R2/2 holey carbon grid. Grids were hand blotted from behind using Whatman paper #1 and plunged into liquid ethane using a Mark IV Vitrobot (Thermo Fisher Scientific) for vitrification. Samples were then transferred under liquid nitrogen into a cryo side-entry holder (Gatan) and loaded into a JEM 2100 cryo TEM (JEOL) operating at 200 k. Holes containing vesicles were targeted for data collection. A nominal magnification of 40,000x was used, which corresponded to a pixel size of 0.29 nm/pixel on our 4k ultrascan CCD (Gatan).

### Statistics

Data are presented as mean ± standard derivation of the mean (SD). Statistical analysis was performed by using Prism (GraphPad, Inc., La Jolla, CA, USA). Western blots were analyzed using Image J software. Two-way analysis of variance was used to determine differences among groups. When identified, a student’s t-test was used to compare differences between groups. A p-value of < 0.05 was considered statistically significant.

## RESULTS

### Expression of R345W-Fibulin-3 alters EV size in RPE cells

We first optimized conventional filtration and ultracentrifugation as well as density gradient centrifugation to isolate EVs from cell culture media of RPE cells. Conventional filtration and ultracentrifugation (P100K) and density gradient centrifugation (Pgrad) were used to isolate EVs. TEM and Cryo-EM were performed to study the morphology of the EVs (Pgrad). TEM imaging showed concave-appearing vesicles (**Fig. 1 A**) and Cryo-EM showed spherical vesicles (**Fig. 1 B**) both with two subpopulations of EVs: a small group with diameters around 30nm and a large group with diameters around 100nm. Moreover, TEM showed an increased amount of small EVs (~30 nm) in the mutant group compared to the WT group. This result was further confirmed by NTA showing that, in the mutant group, the particle size distributions were smaller than those of the WT EVs **(Fig. 1 C**). There were no significant differences in the amount of protein per vesicle between WT and mutant groups or in the total amount of particles in these two groups (**Fig. 1 D**). Western immunoblots confirmed the presence of EV markers (P100K), including EGFR, Flotillin-1, Alix, and Gluc-tagged Fibulin-3 are expressed in both WT and mutant EVs (**Fig. 1 E**).

**Figure 1.**
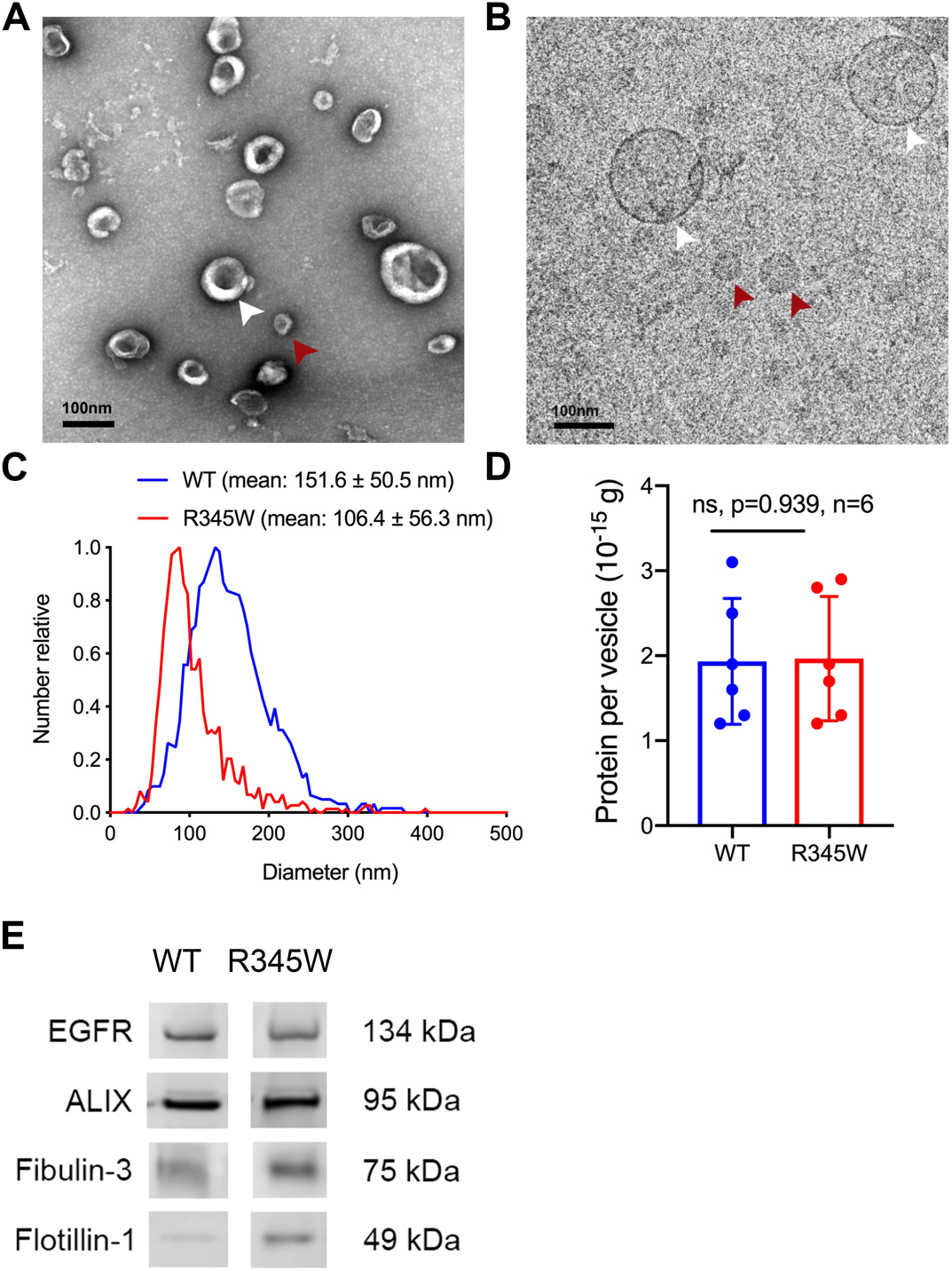
Expression of R345W-Fibulin-3 in RPE cells alters EV size. (A-B) TEM of EVs reveals concave structure and two subpopulations: small vesicles ~30 nm in diameter (red arrowhead) and large vesicles ~100nm in diameter (white arrowhead). Scale bar: 200 nm. (C) Nanoparticle tracking analysis confirms a median diameter of 94.5 nm and 143.5 nm for mutant and WT EVs, respectively. (D) No significant differences in the amount of protein per vesicle or in the total amount of vesicles between WT and mutant groups are observed. (E) Western immunoblots confirmed the presence of EV markers, including EGFR, Flotillin-1, Alix, and Fibulin-3 in both WT and mutant EVs.

### Expression of R345W-Fibulin-3 in RPE cells partially alters their EV cargo content

EVs derived from WT and mutant ARPE-19 cells were purified using sucrose cushion and iodixanol buoyant density gradient ultracentrifugation (Pgrad). Final ultracentrifugation yields 10 remaining fractions, with EVs in the third fraction from the top (density=1.096) (Choi and Gho, 2015). SDS-PAGE gels showed distinct differences in protein distribution pattern between these groups (**Supplemental Fig. 1 A**). Western immunoblots confirmed that the presence of EV markers (Pgrad), including Alix, HSP70, Flotillin-1 and Gluc-tagged Fibulin-3, in the EV fraction (**Supplemental Fig. 1 B**).

EVs (Pgrad) derived from WT and mutant ARPE-19 cells were subjected to proteomic analysis. A total of 2,906 proteins were identified in WT-Fibulin-3 EVs, and 2,899 proteins were identified in mutant Fibulin-3 EVs. Pathway analysis revealed that WT ARPE-19 cells secrete EVs containing components of the primary cilia and sonic hedgehog signaling (SHH) pathways, suggesting a role for RPE EVs in the maintenance of RPE cell and photoreceptor differentiation (Wan et al., 2007;Amirpour et al., 2012). In contrast, mutant ARPE-19 cells were found to secrete vesicles containing EMT drivers and components of the lysosomal degradation pathway, indicating that EVs derived from R345W-Fibulin-3 RPE cells potentially contribute to EMT (**Fig. 2 A**). The log differential expression (Log2DE) was calculated by determining the differential expression of R345W and WT, then taking the log base 2 of that value. Downregulated proteins in the SHH and primary cilia pathways and upregulated EMT markers are listed in the table (**Fig. 2 B**).

**Figure 2.**
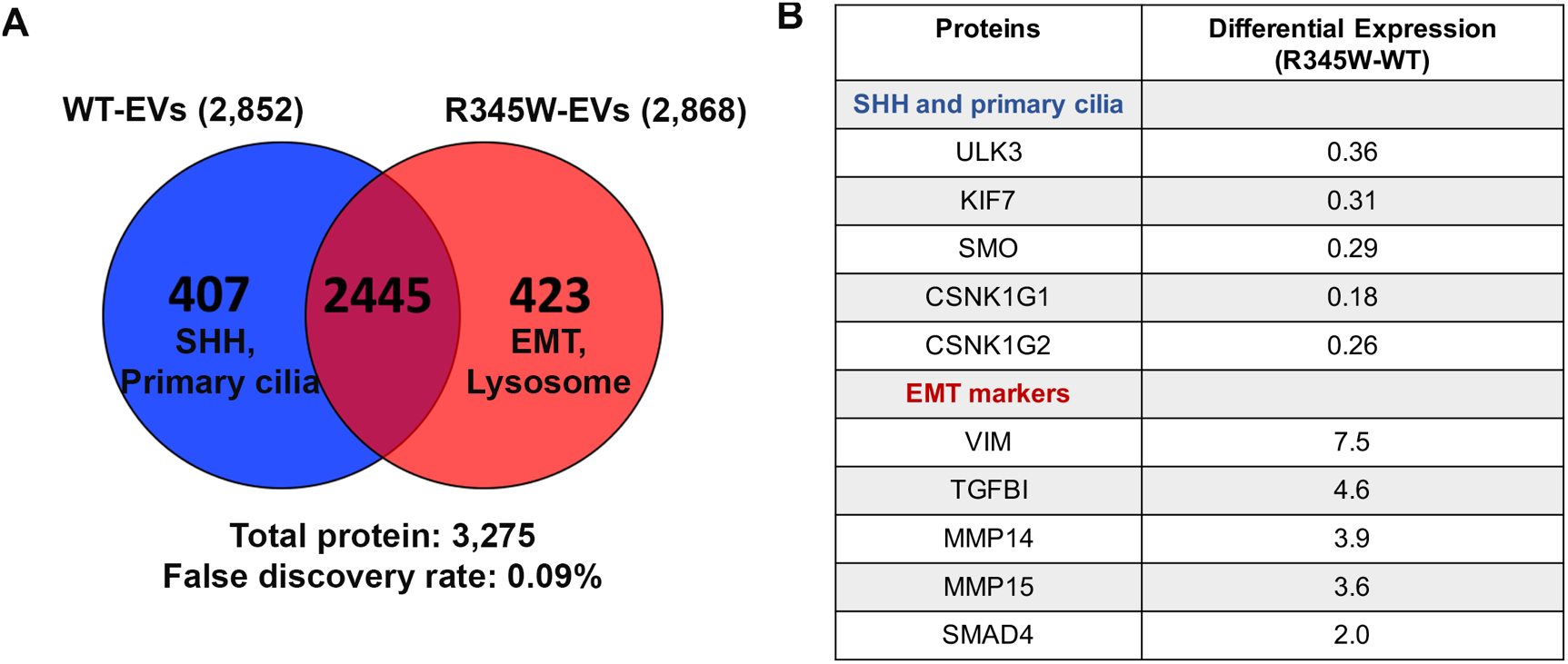
Proteomic analysis of RPE EVs reveal markers of EMT. (A) A total of 2,906 proteins were identified in WT-Fibulin-3 EVs, and 2,899 proteins were identified in R345W-Fibulin-3 EVs. Pathway analysis revealed that ARPE-19 cells overexpressing WT-Fibulin-3 secrete EVs containing components of the sonic hedgehog signaling pathway and primary cilia. In contrast, ARPE-19 cells expressing R345W-Fibulin-3 secreted vesicles containing components of EMT markers and lysosome components. (B) Table of selected proteins and fold-change in spectral counts from proteomic profiling between WT RPE EVs and mutant RPE EVs.

### R345W-RPE-derived EVs have increased TGF-β1 protein

TGF-β is a well-known regulator of EMT (Xu et al., 2009). Recent studies have shown that EV-bound TGF-β1 can induce EMT in recipient cells (van de Vlekkert et al., 2019). To test whether induction of mutant Fibulin-3 alters TGF-β protein content, we quantified TGF-β1 protein in cell culture media, supernatant, EVs in both WT and mutant groups using a commercial ELISA. Cell culture media were collected from ARPE-19 cells expressing either WT-Fibulin-3 or R345W-Fibulin-3 after culturing for 48 hours. EVs (P100K) were isolated from WT and mutant ARPE-19 cells and supernatant was collected as well. Media left over after EVs are centrifuged out were collected as supernatant. Our experimental results revealed that compared to the WT EVs, TGF-β1 protein was significantly more abundant in the mutant EVs (each experiment was performed in duplicate, n=3, p<0.01) (**Fig. 3A**). In addition, compared to the WT media and supernatant, the amount of TGF-β1 protein was significantly greater in the mutant media and supernatant (each experiment was performed in duplicate, n=3, p<0.01) (**Fig. 3B**). These results suggest that expression of mutant Fibulin-3 induces a higher abundance of TGF-β1 protein in RPE cells and EVs derived from RPE cells.

**Figure 3.**
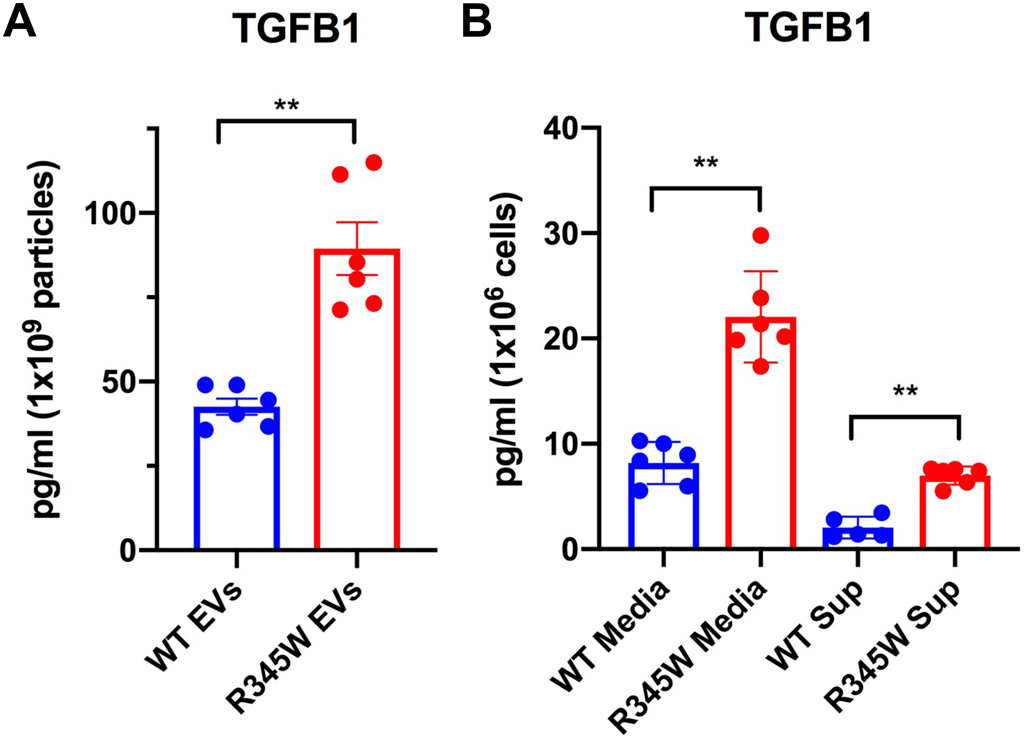
Mutant EVs demonstrate increased TGF-β1 activity. TGF-β1 activities in media, supernatant, and EVs from WT and mutant groups were quantified by ELISA. Mutant EVs show significantly greater TGF-β1 protein compared to WT EVs (each experiment was performed in duplicate, n=3, p<0.01). In addition, mutant media and supernatant contain significantly more TGF-β1protein (each experiment was performed in duplicate, n=3, p<0.01).

### R345W-RPE-derived EVs are sufficient to induce EMT in recipient RPE cells

EV transfer experiments were conducted to determine if EVs from mutant cells are sufficient to promote EMT in uninfected cells. EVs (P100K) were isolated from ARPE-19 cells expressing WT-Fibulin-3 or R345W-Fibulin-3. Control (uninfected) ARPE-19 cells were seeded onto a 96-well plate (5×10^4^/well). On Day 1, the wound was created, each well was rinsed with PBS, and media containing WT or mutant EVs were added to the appropriate wells. A total of n=8 was used for each group, and the experiment was repeated three times with new EV preparations. EV concentrations were chosen according to a previously published study (van de Vlekkert et al., 2019). For these studies, 1 x10^10^/mL EVs were used. A schematic for the scratch assay timeline is shown (**Fig. 4**). A representative experiment is shown below (**Fig. 5A**). To compare across groups, the maximum recovery at 72 hours was used to determine the time required to close half of the wound area. The mean half-max wound recovery time across three experiments revealed that EVs from mutant cells had the fastest wound recovery time (R345W= 17hrs, WT = 24hrs, No EVs = 21hrs (**Fig. 5B**)).

**Figure 4.**
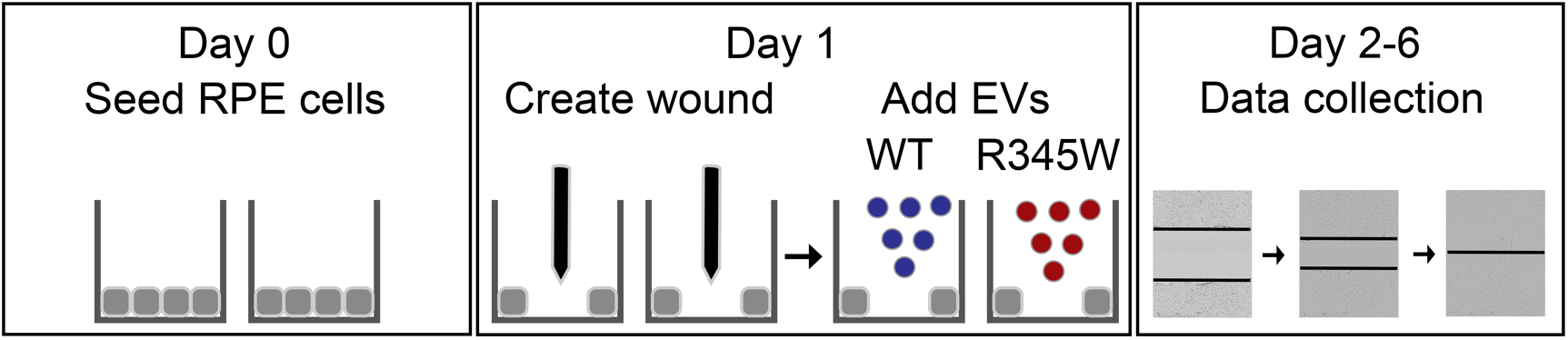
Schematic of EV transplant experiments. On Day 0, control ARPE-19 cells were plated onto a 96-well plate. On Day 1, the wound was created, each well was rinsed with PBS, and media containing WT or mutant EVs were added to the appropriate wells. On Day 3-6, recovery rate was automatically analyzed by the software.

**Figure 5.**
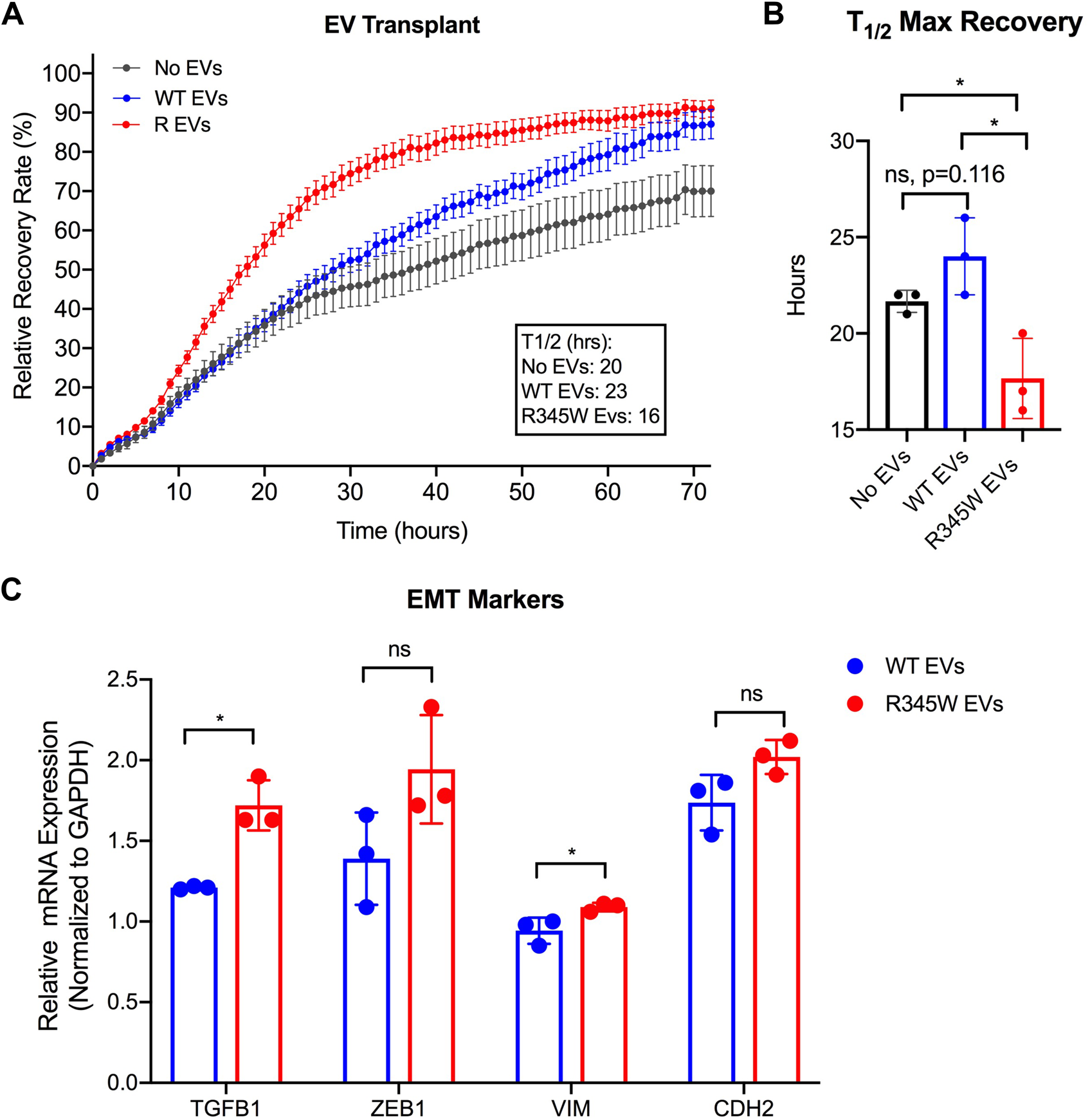
RPE-derived EVs are sufficient to induce EMT in RPE cells. (A-B) Scratch assays show that compared to the cell-only (T1/2=21± 0.5) and cells exposed to WT EVs groups (T1/2=24± 1.6), exposing ARPE-19 cells to mutant RPE cell-derived EVs resulted in a faster rate of 50% wound closure (T1/2=17± 1.7). (C) The relative quantity of mRNA in uninfected ARPE-19 cells was normalized to GAPDH and then normalized to the control group (No EVs). RT-PCR analysis show that compared to the control (No EVs), the mRNA expression levels of four EMT markers were significantly greater in cells treated with mutant EVs (n=3, p<0.01). Compared to WT EVs, mutant EVs increased the expression of TGF-β1 and VIM mRNA (n=3, p<0.01).

**Figure 6.**
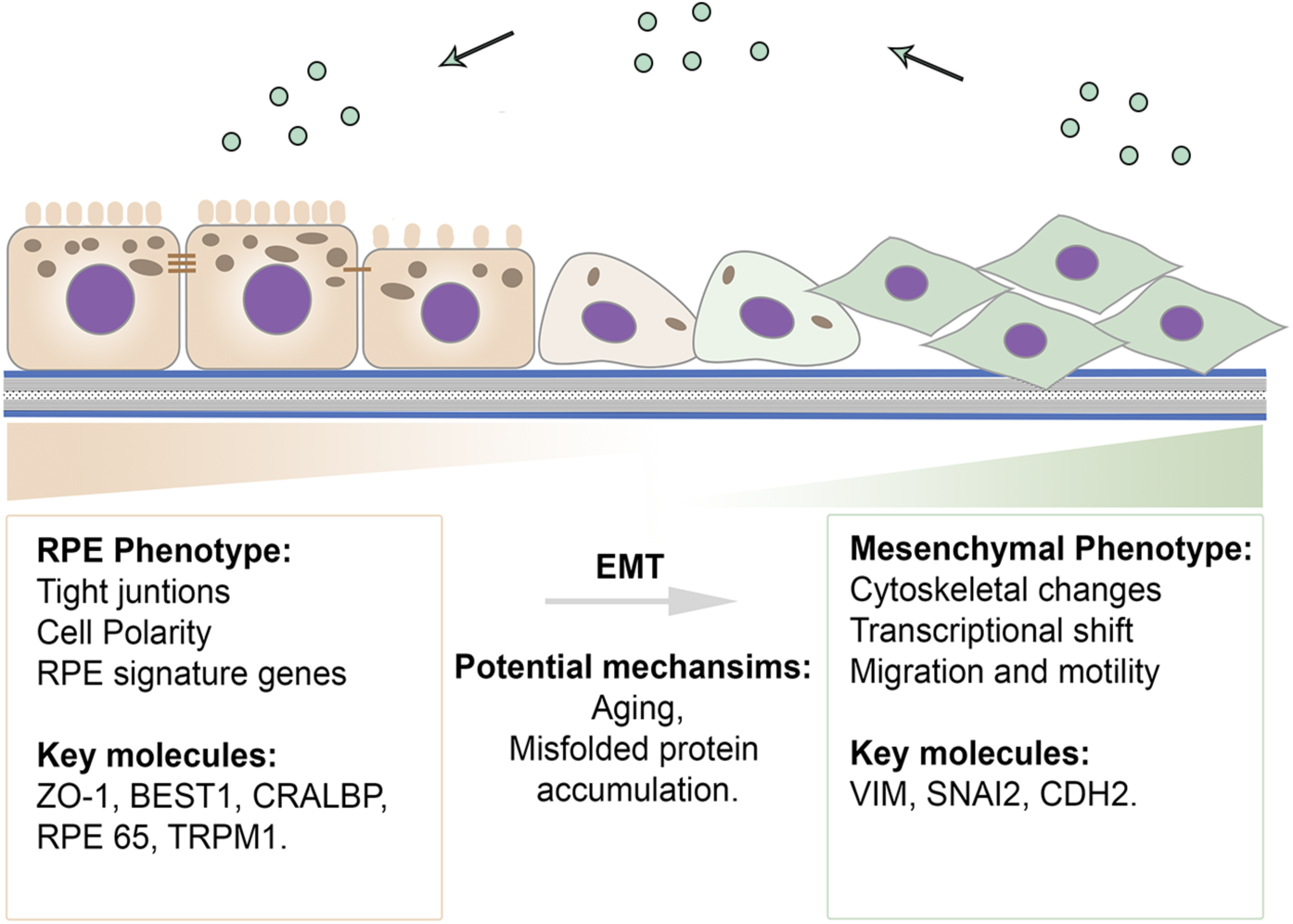
Schematic illustration of proposed scheme. RPE EVs act in a paracrine manner to impact adjacent RPE cell differentiation. The green circles represent EVs derived from RPE cells expressing R345W-Fibulin-3.

Using RT-PCR, we compared the expression levels of EMT-promoting factors (TGF-β1, (Zinc Finger E-Box Binding Homeobox 1 (ZEB1), vimentin (VIM), and Cadherin 2 (CDH2)) in each group 24 hours after EV treatment. The relative quantity of mRNA was normalized to GAPDH and then normalized to the control group (No EVs). Compared to the control group (No EVs), the mRNA expression levels of four EMT markers were significantly greater in cells treated with mutant EVs (n=3, p<0.01). Compared to WT EVs, mutant EVs increased the amount of TGF-β1 protein and VIM (n=3, p<0.01). Taken together, these data suggest a role for TGF-β1 in EV-induced EMT of ARPE-19 cells (**Fig. 5C**).

## Discussion

Under normal conditions, RPE cells are terminally differentiated cells that rarely divide or proliferate (Kozlowski, 2012). Recent studies in post-mortem age-related macular degeneration (AMD) eyes have found upregulation of critical EMT drivers such as TGF-β, VIM, and Snail (Hirasawa et al., 2011;Ishikawa et al., 2016;Ghosh et al., 2018). These findings suggest that dysfunctional RPE cells are capable of migration and proliferation. In other tissues, EMT is regulated, in part, by EVs, suggesting that this mechanism may also be possible in RPE cells(Vella, 2014;Kim et al., 2016;Chen et al., 2017).

In this study, we investigate the role of EVs in the induction of EMT in RPE cells. We first showed that expression of the misfolded protein R345W-Fibulin-3 alters the size of EVs. TEM showed concave-appearing vesicles while cryo-EM imaging showed spherical vesicles. In both forms of imaging there were a variety of different sizes which segregated as “small” (about 30 nm diameter) and “large” (over 100 nm diameter) using NTA, confirming that EVs derived from mutant cells were smaller on average than those derived from WT cells. Despite this difference in vesicle size the amount of protein was not significantly different between the two vesicle types, suggesting that the size difference alone between these two populations does not have a broad impact on the amount of vesicle cargo. A recent study demonstrated, however, that the complexities of heterogeneous EV subpopulations may imply distinct biological functions (Zhang et al., 2018), further emphasizing the importance of separating and studying subpopulations of EVs.

We next examined whether expression of R345W-Fibulin-3 alters EV cargo. Interestingly, we found that EVs derived from RPE cells overexpressing WT-Fibulin-3 contain critical members of SHH and primary cilium pathways. SHH coordinates photoreceptor differentiation during development (Stenkamp et al., 2000) and the primary cilium pathway regulates cellular differentiation and division by mediating hedgehog and Wnt signaling.(Oh and Katsanis, 2012). Prior studies have shown that abnormal activation or absence of primary ciliary signaling is associated with less differentiation and uncontrolled cell division (Spalluto et al., 2013). Diminished SHH and ciliary tip signaling in EVs derived from RPE cells overexpressing R345W-Fibulin-3 suggests that the RPE cells may lose their capacity to provide effective support mechanisms that maintain photoreceptor outer segments.

TGF-β ELISAs revealed greater TGF-β1protein content in mutant EVs, consistent with the proteomic data showing that EMT drivers were found to be 3-to 7-fold more abundant in R345W EVs compared to WT EVs. This finding is consistent with our prior work showing that expression of R345W-Fibulin-3 activates the UPR in RPE cells. Specifically, the IRE1/XBP1 pathway is preferentially upregulated after expression of R345W-Fibulin-3 in primary RPE cells (Zhou et al., 2020b). The UPR and TGF-β-induced EMT signaling pathways interact in an IRE1-dependent manner (Urano et al., 2000;Santibanez, 2006;Liu et al., 2019). Taken together, these data suggest a role for TGF-β1 in EV-induced EMT of ARPE-19 cells.

EV transplant experiments were conducted to determine if EVs from mutant cells are sufficient to promote EMT. Previous studies in our lab showed that expression of mutant Fibulin-3 accelerates would healing (Zhou et al., 2020b). In this study, our experiments reveal that mutant EVs accelerate RPE proliferation and migration to a similar extent as does the expression of mutant Fibulin-3, suggesting that EVs play a major role in driving EMT of RPE cells. Moreover, the data show that expression levels of EMT-promoting factors were greater in the after mutant EV treatment. Taken together, our data showed that mutant EVs are sufficient to induce EMT in ARPE-19 cells, suggesting that RPE-derived EVs act in a paracrine manner to influence adjacent RPE cell differentiation.

## Conclusion

In summary, our experimental results indicate that misfolded proteins in RPE cells alter specific EV cargo, resulting in EVs that are sufficient to induce EMT of RPE cells. **Figure 6** illustrates our working model for EV-induced RPE EMT. The findings from this study will help elucidate the mechanisms underlying RPE dysfunction in inherited macular degenerations.

---

**Supplemental Figure 1. Expression of R345W-Fibulin-3 in RPE cells alters EV content**. (A) EVs derived from ARPE-19 cells overexpressing WT-Fibulin-3 or R345W-Fibulin-3 were purified using sucrose cushion and iodixanol buoyant density gradient ultracentrifugation. SDS-PAGE gels showed distinct differences in protein distribution patterns between these groups. WT cell lysates (WCL) and cell culture media (CCM) were used as controls. (B) Western immunoblots confirmed the presence of EV markers in EVs isolated from the density gradient ultracentrifugation, including Alix, HSP70, Flotillin-1 and Gluc-tagged Fibulin-3.

