## Supplementary figures and images for "Extracellular vesicles from retinal pigment epithelial cells expressing R345W-Fibulin-3 induce epithelial-mesenchymal transition in recipient cells"

### Supplemental Figure 1

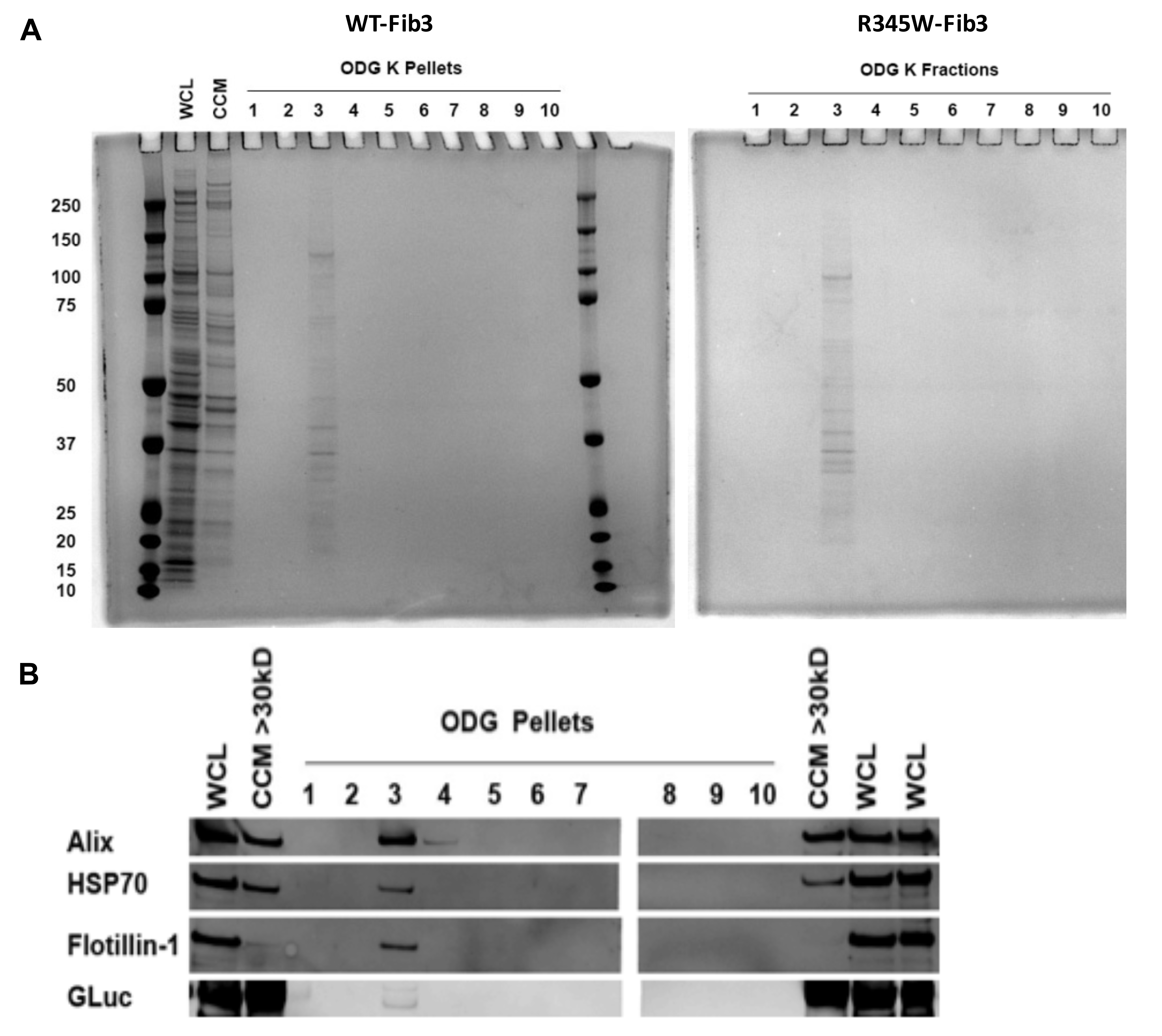
